# Neurovascular coupling preserved in a chronic mouse model of Alzheimer’s disease: Methodology is critical

**DOI:** 10.1101/474916

**Authors:** Paul S Sharp, Kam Ameen-Ali, Luke Boorman, Sam Harris, Stephen Wharton, Clare Howarth, Peter Redgrave, Jason Berwick

**Author notes:** **Corresponding author**: Dr Jason Berwick, Department of Psychology, University of Sheffield, Floor D, Cathedral Court, 1 Vicar Lane, Sheffield, South Yorkshire, UK, S11HD.

## Abstract

Neurovascular coupling is the process by which neural activity causes localised changes in cerebral blood flow. Impaired neurovascular coupling has been suggested as an early pathogenic factor in Alzheimer’s disease (AD), and if so, could serve as an early biomarker of cerebral pathology. We have established an anaesthetic regime in which evoked hemodynamic responses are comparable to those in awake mice. This protocol was adapted to allow repeated measurements of neurovascular function over three months in the hAPP-J20 mouse model of AD (J20-AD) and wild-type (WT) controls. Animals were 9-12 months old at the start of the experiment, which is when deficits due to the disease condition would be expected. Mice were chronically prepared with a cranial window through which optical imaging spectroscopy (OIS) was used to generate functional maps of the cerebral blood volume and saturation changes evoked by whisker stimulation and vascular reactivity challenges. Unexpectedly, the hemodynamic responses were largely preserved in the J20-AD group. This result failed to confirm previous investigations using the J20-AD model. However, a final acute electrophysiology and OIS experiment was performed to measure both neural and hemodynamic responses concurrently. In this experiment, previously reported deficits in neurovascular coupling in the J20-AD model were observed. This suggests that J20-AD mice may be more susceptible to the physiologically stressing conditions of an acute experimental procedure compared to WT animals. These results therefore highlight the importance of experimental procedure when determining the characteristics of animal models of human disease.

**Significance Statement:** Using a chronic anaesthetised preparation, we measured hemodynamic responses evoked by sensory stimulation and respiratory gases in the J20-AD mouse model of Alzheimer’s Disease over a period of 3 months. We showed that neurovascular responses were preserved compared to age matched wildtype controls. These results failed to confirm previous investigations reporting a marked reduction of neurovascular coupling in the J20-AD mouse model. However, when our procedure involved acute surgical procedures, previously reported neurovascular deficits were observed. The effects of acute electrode implantation were caused by disturbances to baseline physiology rather than a consequence of the disease condition. These results highlight the importance of experimental procedure when determining the characteristics of animal models of human disease.

## Introduction

Alzheimer’s disease (AD) in humans currently has no effective treatment. Most of our recent insights into the causes of AD are linked to the discovery of increased amounts of Beta-Amyloid plaques associated with significant neuronal loss(Hardy, 1992). Despite the large investment in both clinical and pre-clinical research resources focussed on the Beta-Amyloid hypothesis, no effective preventative or remedial treatment has so far been found. Consequently, alternative theories regarding the onset and development of AD have been proposed. One of the notable rival hypotheses is the Neurovascular Degeneration Hypothesis (NDH), first proposed by Zlokovic (Zlokovic, 2010, 2011). It suggests that a functional deficit within cells of the neurovascular unit (neurons, glial cells, pericytes and vascular cells) would deprive active neurones of adequate oxygen and glucose, which could either be the initial trigger of AD, or significantly add to the disease burden. If correct, the NDH would offer the potential for developing new treatments based on targeting identified dysfunctional cells in the neurovascular unit. With the aid of modern neuroimaging methods (e.g. functional magnetic resonance imaging – fMRI) measurements of the progressive breakdown of neurovascular coupling (NVC) could also act as an early biomarker of AD. A recent study analysing data from the Alzheimer’s Disease Neuroimaging Initiative (ADNI, (Iturria-Medina et al., 2016)) suggested that cerebrovascular dysfunction may be the earliest pathological event to emerge, possibly marking the beginning of the disease process. This report alone highlights the need for more research into the NDH.

A problem facing investigations of the neurovascular unit is that the technologies currently used with human subjects do not have the spatial and temporal resolution required to understand the basic mechanistic processes involved. To address this issue transgenic mouse models of AD have been developed and now play an important role in investigating the underlying mechanisms responsible for the AD-like disease state. These transgenic mouse models may not perfectly replicate the human condition, however, it is hoped that they share the critical features to enable us to discover the underlying mechanisms responsible for the condition (Ameen-Ali et al., 2017). The pre-clinical AD mouse models allow the use of more invasive technologies to measure both hemodynamic and neuronal variables in more detail throughout disease development. Consequently, several groups have reported disruptions to NVC in transgenic AD mice that overexpress human amyloid precursor protein (hAPP) (Lacoste et al., 2013, Ongali et al., 2014, Park et al., 2014, Royea et al., 2017). These studies used laser Doppler flowmetry (LDF) to measure cerebral blood flow (CBF) from a restricted cortical region in acutely anaesthetised animals. To extend these investigations we have developed a chronic mouse preparation that is sufficiently stable to permit repeated measures of neurovascular function across a 3 month period when the disease state is present (Sharp et al., 2015). Consequently, with the J20-AD mice and age matched wild type controls temporarily anaesthetised, we used repeated measures 2-dimensional optical imaging spectroscopy (2D-OIS) to record neurovascular responses evoked by sensory stimulation and vascular reactivity challenges. A final acute experiment was conducted in which 2D-OIS and multi-channel electrophysiology were performed simultaneously. Contrary to expectation, our chronic imaging studies found little or no difference in the hemodynamic responses between J20-AD and WT animals across a range of sensory stimulations and gas challenges. However, in the final acute experiment, reduced NVC function similar to that reported in previous investigations (Lacoste et al., 2013, Ongali et al., 2014, Park et al., 2014, Royea et al., 2017) was observed. This difference suggests that the J20-AD mice are more susceptible to the physiological stresses incurred by acute experimental procedures compared with WT controls. These results indicate that experimental conditions are critical when characterising mouse models of human disease, and differences in methodology are likely responsible for the variable and sometimes incompatible results frequently seen in mouse pre-clinical neurovascular studies.

## Methods

### Anesthesia and cranial window surgery

All animal procedures were performed in accordance with the guidelines and regulations of the UK Government, Animals (Scientific Procedures) Act 1986, the European directive 2010/63/EU, and approved by the University of Sheffield Ethical review and licensing committee. Two groups of 5 animals were used: (i) heterozygous transgenic male C57BL/6 mice that overexpress human APP (hAPP) carrying the human Swedish and Indiana familial AD mutations under control of the PDGFβ chain promoter (hAPP-J20 line); and (ii) male wild type controls. All animals were between 9 and 12 months on the date of surgical preparation. Animals were anesthetized with fentanyl-fluanisone (Hypnorm, Vetapharm Ltd), midazolam (Hypnovel, Roche Ltd) and sterile water (1:1:2 by volume; 7.0 ml/kg,i.p.) for surgery, while anaesthesia during the imaging experiments was further maintained with isoflurane (0.5–0.8%) in 100% oxygen. A homoeothermic blanket (Harvard Apparatus) maintained rectal temperature at 37°C. Mice were placed in a stereotaxic frame (Kopf Instruments) and a dental drill was used to thin the skull overlying the right somatosensory cortex to translucency thereby forming an optical window (~4mm^2^). A thin layer of clear cyanoacrylate cement was applied to reinforce and smooth the window. This reduced specular reflections during imaging and prevented skull regrowth during the 3-month experimental period. A stainless steel imaging chamber was secured over the thinned cranial window using dental cement. This chamber was used to stabilise the head during imaging sessions. After surgery animals were left to recover for a minimum of 7 days before chronic experimental imaging sessions commenced. During this period animals were monitored regularly and weighed to ensure they did not fall below 90% of surgical day body weight.

### Experimental Design

All animals were imaged using an anaesthetised condition adapted from our previous published methodology (Sharp et al., 2015). All animals underwent 4 imaging sessions. The first session occurred 7-10 days after cranial window surgery. The following three sessions were each separated by ~30 days. In the last session a 16-channel electrode was placed in the active whisker barrel region (see below for method of localisation) to provide a measure of concurrent hemodynamic and neural activity. Each session began with an induction of anaesthesia (see methods) followed after 1hour by 2D-OIS. In the last acute (4^th^) imaging session imaging started 30 minutes after the electrode had been inserted. Each 2D-OIS session comprised 8 separate experiments performed in a set order. Each experiment separated from the previous one by ~3mins during which time we acquired a dark image used to remove camera noise from the spectroscopy data.

#### Exp1

Short duration sensory stimulation breathing 100% oxygen: With the animal breathing oxygen (100%) there were 30 successive trials of 25s. 5s after the start of each trial whisker stimulation was administered for 2s. Together, 750s of continuous data was collected. For all stimulation experiments the whiskers contralateral to the imaging chamber were mechanically deflected using aplastic T-bar attached to a stepper motor under computer control. Whiskers were deflected ~1 cm in the rostro-caudal direction at 5Hz.

#### Exp2

Exp1 was repeated to ensure preparation and recording stability.

#### Exp3

Mild gas challenge: while the animal transitioned from breathing oxygen to normal air. 2D-OIS lasted 750s with the transition to air occurring after 105s.

#### Exp4

Short duration sensory stimulation breathing air: Used the same paradigm as Exp 1 except the animal was breathing normal air rather than 100% oxygen

#### Exp5

Long duration sensory stimulation breathing air: 15 x 70s consecutive trials with 16s whisker stimulation (5Hz) presented after 10s. Total record comprised 1050s of data.

#### Exp6

Mild gas challenge: while the animal transitioned from breathing air back to 100% oxygen. 750s of data were recorded with the transition occurring after 105s

#### Exp7

Long duration sensory stimulation under 100% oxygen: 15 x 70s consecutive trials, with 16s whisker stimulation at 5Hz occurring after 10s. Total record comprised 1050s of data.

#### Exp8

Major gas challenge in the form of hypercapnia: 250s after starting the record the animal was switched from breathing 100% oxygen to 90% and10% carbon dioxide. The duration of this challenge was 250s, after which a further 250s of data were collected giving a total of 750s of recorded data.

### Statistical Analysis

The first question we addressed were whether the evoked haemodynamic responses changed over time (sessions 1-3) and were different between the J20-AD and WT groups. Separate 2-way repeated measures ANOVA’s were performed for Hbt, Hbo and Hbr for each of the 5 whisker stimulation experiments (Exp1,2,4,5,7). A single measure, the mean response size from the stimulation period was calculated and used for the analysis. Based on the results of this analysis (see below) a second repeated measures ANOVA was performed. In this analysis we averaged the three chronic imaging sessions together to create a single chronic session for both WT and J20-AD mice with the repeated measurement now being stimulation experiment (Exp1,2,4,5,7). This same analysis was also performed for the acute imaging session. This would show if there were any differences between the WT and J20-AD animals in the chronic or acute parts of the study.

### 2-Dimensional optical imaging spectroscopy (2D-OIS)

2D-OIS was used to estimate changes in cortical oxyhemoglobin (HbO_2_), deoxyhemoglobin (Hbr) and total hemoglobin concentration (Hbt). To generate spatial maps of hemodynamic responses, the cortex was illuminated with 4 wavelengths of light (495nm, 559nm, 575nm and 587nm) using a Lambda DG-4 high-speed galvanometer (Sutter Instrument Company, USA). Remitted light was collected using a Dalsa 1M60 CCD camera at 184 × 184 pixels (resolution ~75 μ m), with a frame rate of the 32 Hz, and synchronised to filter switching, giving an effective frame rate of 8 Hz.

The analysis approach combined the absorption spectra of oxy and deoxy-haemoglobin with Monte-Carlo simulations of photons passing through homogeneous tissue to estimate the mean path-length of photons for each wavelength. Images were then analysed on a pixel-by-pixel basis using a modified Beer-Lambert law, which used the generated mean photon path lengths, to convert the detected attenuation for each wavelength into predicted absorption. The absorption values were then used to generate 2D spatiotemporal image-series of the estimates of the changes in total hemoglobin (Hbt), oxyhemoglobin (HbO2) and deoxyhemoglobin (Hbr) from the baseline values. All experiments started with the animal breathing 100% oxygen, for this condition we assumed a concentration of hemoglobin in tissue at 100μM and saturation set at 70%. We then used the gas transition experiment 3 to assess how these baselines changed when the animal breathed normal air. This was calculated on an individual animal basis. The baseline blood volume and saturation values in air were consistent for both WT and J20-AD mice (Hbt concentration WT=99.3±0.42sem, AD=99.7±0.47sem; Saturation WT=62.6%±0.6, AD = 62.6±0.6). All hemodynamic changes were taken as the fractional changes from these baseline estimations.

### Selection of regions of interest (ROI) for 2D-OIS data

In each imaging session we used the 16s Hbt data from Exp7 to select the whisker region of interest. During the stimulation period all pixels that were 1.5 standard deviations above the pre-stimulus baseline were deemed above threshold and a region was drawn around all these activated pixels (see Figure 1). Arteries and veins were identified within the activated region using principal component analyses applied to the HbT and HbR images respectively (see Fig 8).

**Figure 1:**
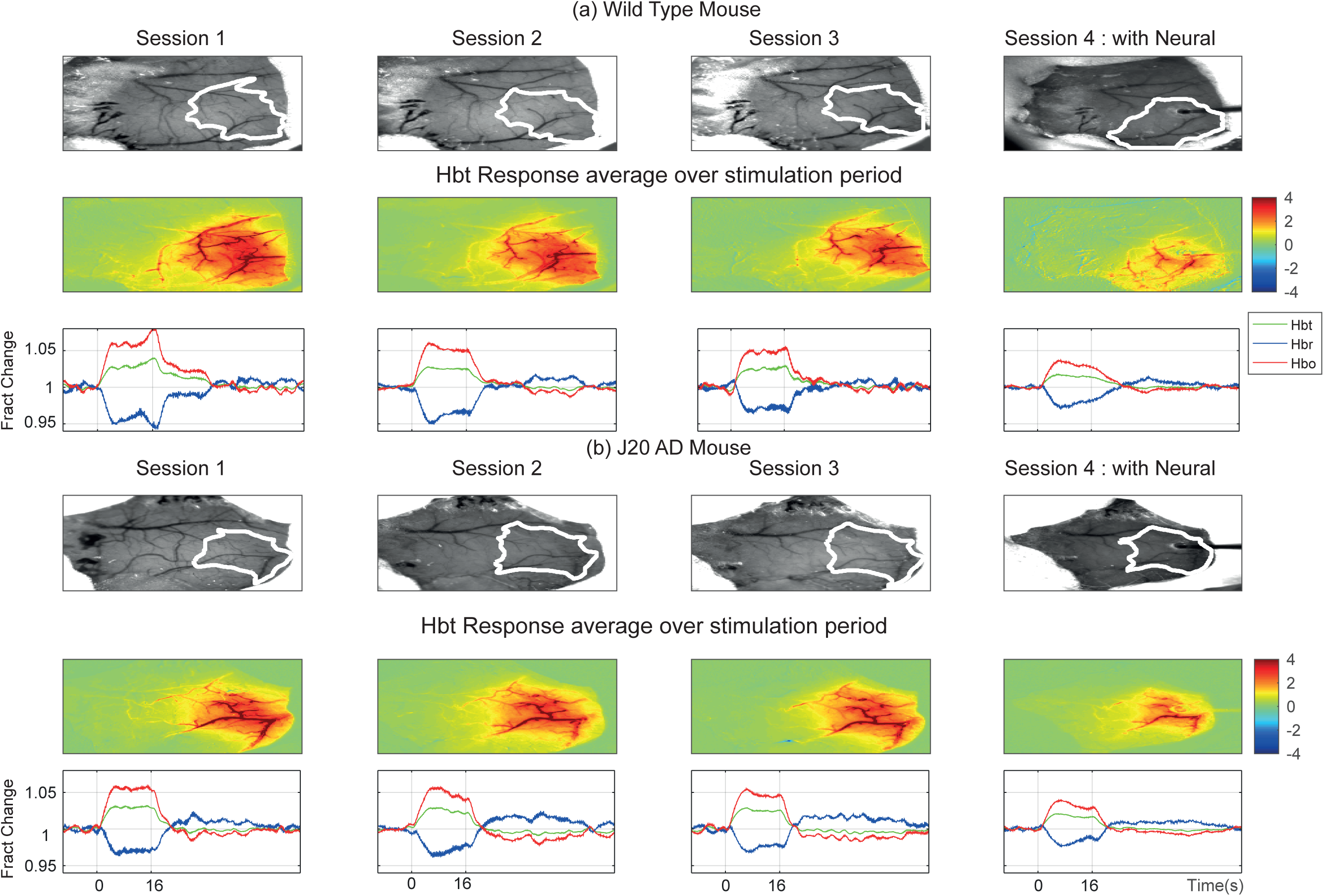
Representative hemodynamic responses from WT (a) and J20-AD mice (b) across three chronic imaging sessions (1-3) each separated by ~30 days and a final acute session (4) where a multi-channel electrode is inserted into the activated whisker region. Activation maps represent the change in Hbt with respect to baseline during a 16s mechanical whisker stimulation. Time series take for Hbt, Hbr and Hbo are taken from the activated whisker region highlighted in white on the reference images (rows 1 and 3). Hbt= total blood volume, Hbo=Oxyhemoglobin, Hbr=deoxyhemoglobin.

**Figure 8:**
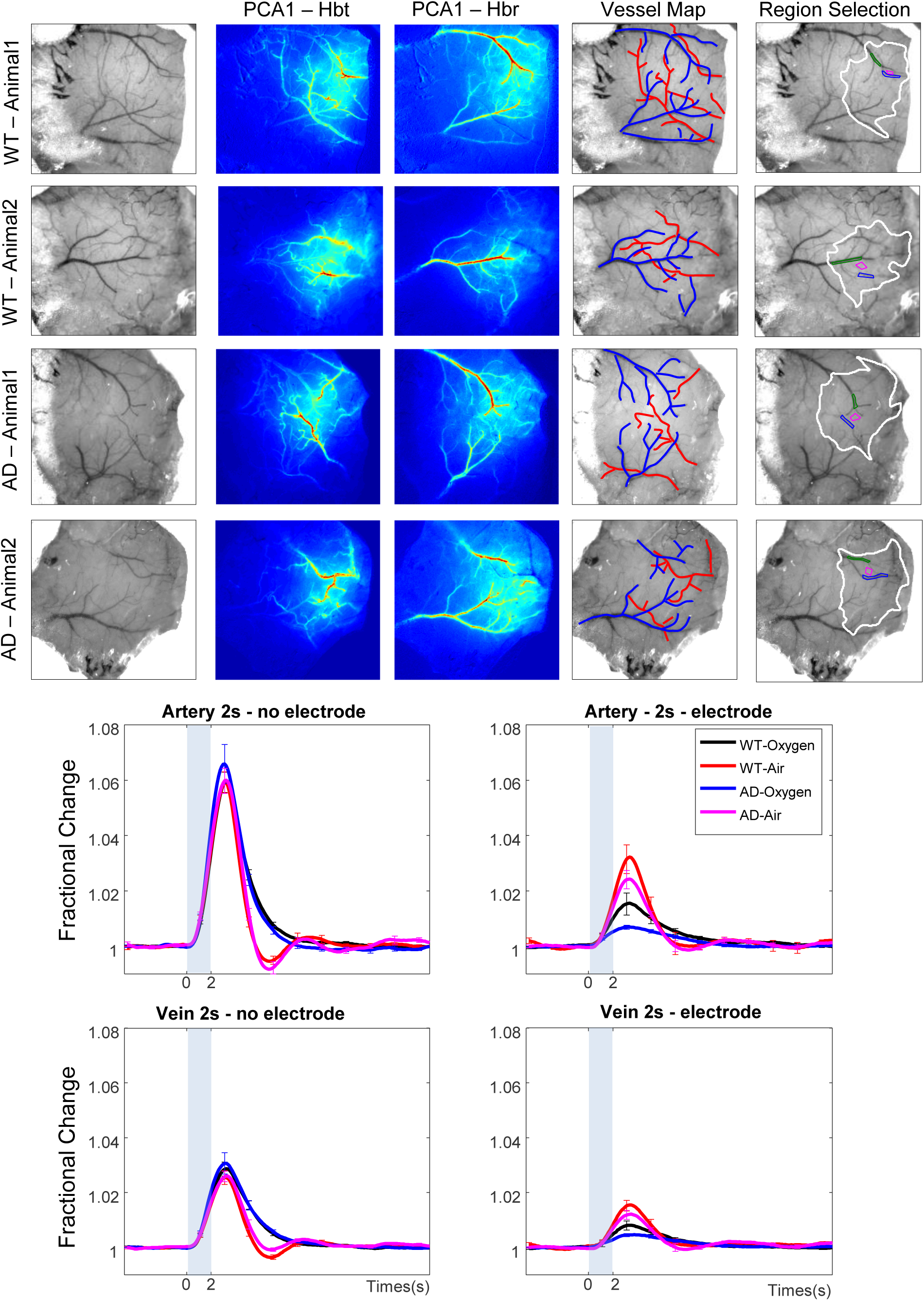
Anatomically compartmentalised vascular responses for 4 representative animals (a) Reference image showing cortical vasculature (b) The first principal component of the Hbt image data block (c) First principal component of the Hbr image data block. (d) Reference images with superimposed surface arteries and veins (e) Calculated whisker response region with selected artery and vein regions. (f) Arterial response time series for chronic 2s stimulation breathing air or oxygen. (g) Arterial response time series for acute 2s stimulation breathing air or oxygen. (h) Venous responses time series for chronic 2s stimulation breathing air or oxygen. (i) Venous response time series for acute 2s stimulation breathing air or oxygen.

### Multi-channel electrophysiology

For the final acute imaging session a 16-channel microelectrode (100μm spacing, site area 177μm^2^, 1.5–2.7 MΩ impedance; Neuronexus Technologies, Ann Arbor, MI, USA) implanted into the right barrel cortex. The microelectrode was positioned to a depth of 1500μm in the centre of the activated whisker region defined previously by 2D-OIS. The electrode was connected to a preamplifier and data acquisition device (Medusa BioAmp/ RZ5, TDT, Alachua, FL, USA). Neural data were sampled at 6 kHz and recorded continuously throughout each of the eight experiments described above.

## Results

### Chronic Imaging: Sensory Stimulation Experiments 1,2,4,5 and 7

The thinned cranial window preparation remained stable throughout the 3 month’s duration of the study with evoked hemodynamic responses remaining remarkably consistent throughout this period for both J20-AD and WT animals (see Figure 1). The first questions we addressed were whether the evoked haemodynamic responses changed over time (sessions 1-3) and were different between the J20-AD and WT groups. Separate 2-way repeated measures ANOVA’s were performed for Hbt, Hbo and Hbr for each of the 5 whisker stimulation experiments (Exp1,2,4,5,7, results shown in Table 1). For each of the three haemodynamic measures (Hbo,Hbr,Hbt), in each of the 5 experiments (Exp1,2,4,5,7) there was only one result that approached significance. For Hbr in Exp7 there was a difference in the sessions (F=5.39 and p=0.01) However, after Bonferroni correction (level needed to be less than p=0.003) this result was also non-significant. All other results showed no statistically significant main effects of session (1-3), or group (WT vs J20-AD). Thus, in all cases differences between the relevant variables failed to reach acceptable levels of statistical reliability (p>0.05, see Table 1). The surprising and remarkable extent to which the haemodynamic responses J20-AD were preserved is illustrated in Figure 2. The stability of response magnitudes across experimental conditions between WT and J20 mice are also summarised in Table 2. Because no reliable differences were found in the results between chronic sessions, we averaged haemodynamic time series across chronic to create sessions for a second analysis to test for differences in the haemodynamic responses within experiments (Exp1,2,4,5 and 7) between the WT and J20-AD groups. For each of the hemodynamic measures (Hbt, Hbr, and Hbo) a 2-way repeated measure ANOVA was conducted which compared responses evoked by the different stimulation conditions and mouse-type (J20-AD vs WT). In each of the analyses the Experiment factor was highly significant (Hbt: F=41.0, p=1.1^10-13^; Hbr: F=21.3, p=1.75^10-9^; Hbo: F=20.6, p=2.8^10-9^), which was to be expected as the stimulations were of varying durations (2s and 16s). On the other hand no reliable statistical differences were found between the two mouse-types for Hbt (F=0.54, p=0.46) and Hbo (F=1.9, p=0.17), however, for Hbr a statistically reliable difference was observed between the groups (F=5.03, P=0.03). This result was due to the washout of Hbr in the J20-AD mice being slightly but consistently smaller than that of the WT animals across experiments (Figure 4). However, again after Bonferroni correction this failed to reach significance (need to be <0.017). In each case the interactions between Experimental condition and mouse-type were also non-significant (Hbt: F=0.05, p=0.99; Hbr: F=0.18, P=0.94; Hbo: F=0.07, p=0.99). Although the experiments within a session were always run in the same order, this time-related confound within these analyses do not obscure the point that the neurovascular performance of J20-AD and WT controls were, for the most part, similar. Specifically, using our chronic procedures we failed to find the large differences reported in previous studies.

**Table 1:** Two way ANOVA to assess changes in Hemodynamic responses between WT and J20-AD across chronic sessions.

|  | Hbt Exp1 | Hbt Exp2 | Hbt Exp4 | Hbt Exp5 | Hbt Exp7 |
| --- | --- | --- | --- | --- | --- |
| Animal Type | F=0.09<br>p=0.77 | F=0.28<br>p=0.60 | F=0.01<br>p=0.94 | F=0.22<br>p=0.64 | F= 0.11<br>p=0.7 |
| Session(1:3) | F=0.18<br>p=0.84 | F=0.01<br>p= 0.99 | F=0.13<br>p= 0.88 | F=1.27<br>p=0.29 | F= 0.42<br>p= 0.66 |
| Interaction | F=0.10<br>p=0.90 | F=1.78<br>p=0.19 | F=1.43<br>p= 0.26 | F=0.20<br>p= 0.81 | F= 0.37<br>p=0.70 |
|  | Hbo Exp1 | Hbo Exp2 | Hbo Exp4 | Hbo Exp5 | Hbo Exp7 |
| Animal Type | F=0.08 p=<br>0.77 | F=0.62 p=<br>0.43 | F=0.14 p=<br>0.71 | F=1.67 p=<br>0.21 | F=0.53 p=<br>0.47 |
| Session(1:3) | F=0.30<br>p=0.74 | F=0.10 p=<br>0.91 | F= 0.36p=<br>0.69 | F= 1.16p=<br>0.33 | F=1.42 p=<br>0.26 |
| Interaction | F=0.06 p=<br>0.94 | F=0.83<br>p=0.45 | F=0.72 p=<br>0.49 | F=0.24 p=<br>0.79 | F= 0.32p=<br>0.73 |
|  | Hbr Exp1 | Hbr Exp2 | Hbr Exp4 | Hbr Exp5 | Hbr Exp7 |
| Animal Type | F=0.05 p=<br>0.822 | F=1.12 p=<br>0.29 | F=1.75 p=<br>0.20 | F= 3.81p=<br>0.06 | F=1.88 p=<br>0.18 |
| Session (1-3) | F=0.58 p=<br>0.56 | F=0.47 p=<br>0.63 | F=1.57 p=<br>0.23 | F=0.43p=<br>0.65 | F=5.39 p=<br>0.01 |
| Interaction | F=0.006 p=<br>0.99 | F=0.02 p=<br>0.97 | F=0.08 p=<br>0.92 | F=0.09 p=<br>0.92 | F=0.45 p=<br>0.64 |

**Table 2:** Average hemodynamic responses (with S.E.M) for chronic and acute session experiments

|  | Average Chronic Session |  |  |  |  | Acute Session (with Electrode) |  |  |  |  |
| --- | --- | --- | --- | --- | --- | --- | --- | --- | --- | --- |
|  | Exp1 | Exp2 | Exp4 | Exp5 | Exp7 | Exp1 | Exp2 | Exp4 | Exp5 | Exp7 |
| WT<br>Hbt | 1.55<br>±<br>0.07 | 1.26<br>±<br>0.06 | 1.40<br>±<br>0.06 | 1.69<br>±<br>0.05 | 1.97<br>±<br>0.04 | 0.56<br>±<br>0.12 | 0.67<br>±<br>0.14 | 0.86<br>±<br>0.14 | 1.23<br>±<br>0.16 | 1.32<br>±<br>0.12 |
| AD<br>Hbt | 1.50<br>±<br>0.12 | 1.20<br>±<br>0.08 | 1.11<br>±<br>0.04 | 1.65<br>±<br>0.06 | 1.93<br>±<br>0.06 | 0.29<br>±<br>0.03 | 0.44<br>±<br>0.04 | 0.67<br>±<br>0.04 | 0.99<br>±<br>0.08 | 0.99<br>±<br>0.13 |
| WT<br>Hbo | 3.02<br>±<br>0.12 | 2.50<br>±<br>0.10 | 2.76<br>±<br>0.10 | 3.27<br>±<br>0.14 | 2.84<br>±<br>0.09 | 1.00<br>±<br>0.25 | 1.31<br>±<br>0.28 | 1.72<br>±<br>0.29 | 2.23<br>±<br>0.24 | 2.34<br>±<br>0.20 |
| AD<br>Hbo | 2.92<br>±<br>0.26 | 2.33<br>±<br>0.19 | 2.62<br>±<br>0.22 | 3.07<br>±<br>0.07 | 3.4<br>±<br>0.11 | 0.39<br>±<br>0.05 | 0.75<br>±<br>0.08 | 1.23<br>±<br>0.10 | 1.67<br>±<br>0.16 | 1.62<br>±<br>0.23 |
| WT<br>Hbr | -1.89<br>±<br>0.07 | -1.63<br>±<br>0.04 | -0.98<br>±<br>0.08 | -0.94<br>±<br>0.07 | -1.73<br>±<br>0.07 | -0.49<br>±<br>0.19 | -0.83<br>±<br>0.2 | -0.71<br>±<br>0.15 | -0.61<br>±<br>0.08 | -1.07<br>±<br>0.18 |
| AD<br>Hbr | -1.82<br>±<br>0.23 | -1.41<br>±<br>0.18 | -0.79<br>±<br>0.11 | -0.64<br>±<br>0.15 | -1.50<br>±<br>0.15 | 0.08<br>±<br>0.03 | -0.31<br>±<br>0.06 | -0.42<br>±<br>0.13 | -0.32<br>±<br>0.14 | -0.46<br>±<br>0.17 |

**Figure 2:**
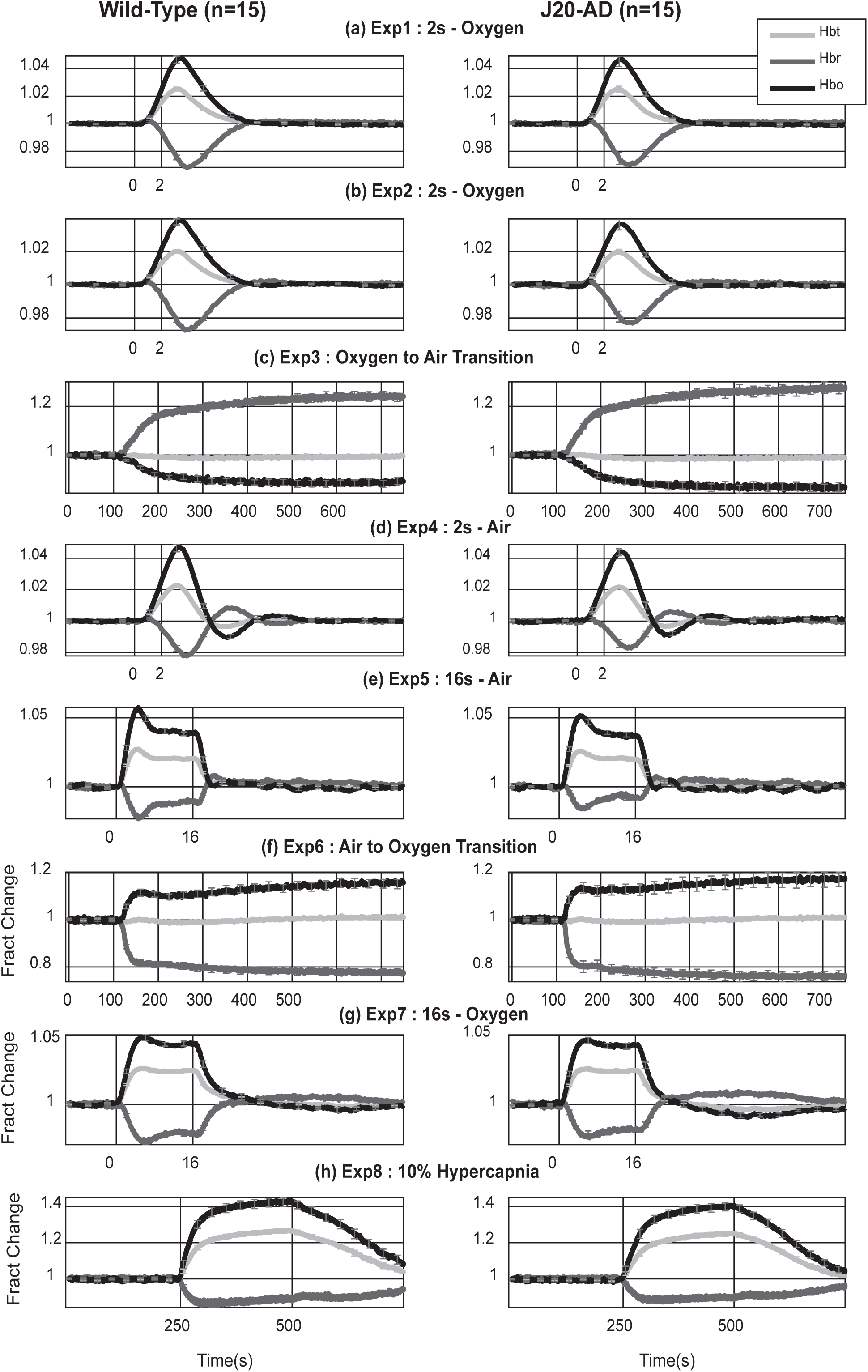
Neurovascular coupling preserved in J20-AD mice. Hemodynamic time series (Hbt, Hbr and Hbo) averaged across chronic sessions for each of the 8 experiments (a-h). WT responses are illustrated in the left column and J20-AD in the right column.

**Figure 4:**
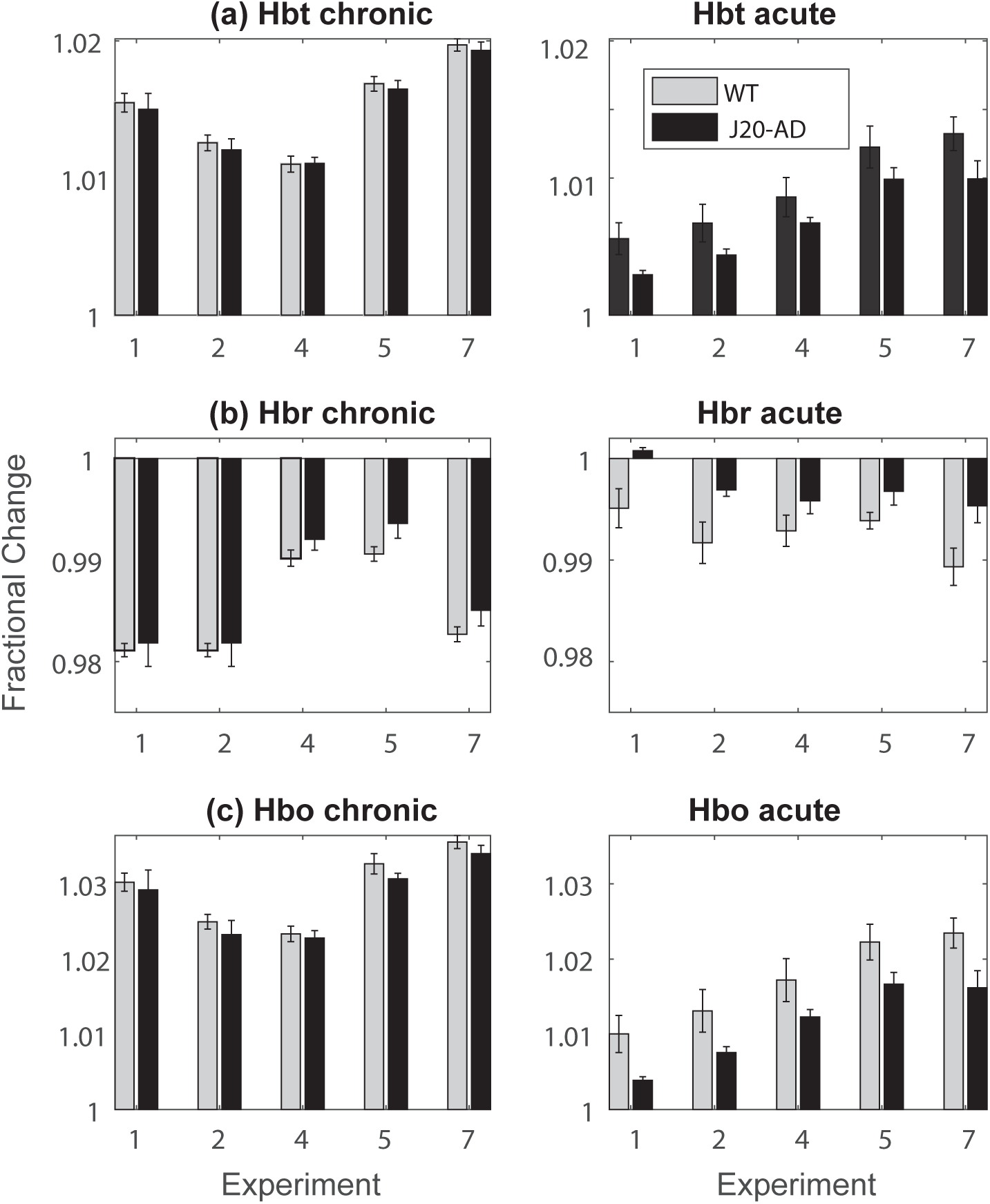
Average magnitude of fractional hemodynamic response for both chronic (left hand side) and acute session (right hand side) stimulation experiments. (a) Hbt (b) Hbr and (c) Hbo.

### Stimulation Experiments 1,2,4,5 and 7 – acute imaging sessions

The 4^th^ and final session involved acute surgical procedures in which a multi-channel recording electrode was implanted prior to the barrel region being imaged. The concurrent acute recording of electrophysiological and hemodynamic variables in this session produced results that were markedly different from previous chronic imaging sessions (See Figures 3 and 4). First, both WT and J20-AD fractional time series responses were reduced compared to the chronic session data. However, the responses of J20-AD mice were dramatically suppressed compared with the corresponding WT responses, especially for experiment 1 (See Figure 3, 4 and Table 2 for overall values). When the same repeated measures ANOVA analyses for Hbt, Hbo and Hbr were performed on data from the acute imaging session, for Hbt there was now a statistically significant difference between the WT and J20-AD groups (F=10.35, p=0.002, level needed after Bonferroni correction 0.017). Differences between the experimental conditions remained significant (F=14.1, p=0.28^10-6^) and there was no reliable interaction between group and experimental condition (F=0.09, p=0.99). This pattern of results was replicated for both Hbo (group, F=16.8, p=0.0002, experimental condition, F=12.3, p= 0.013^10-7^ and interaction, F=0.08, p=0.98), and Hbr (group, F=19.67, p=0.7^10-6^, experimental condition, F=3.1, p= 0.02 and interaction, F=0.42, p=0.79). Thus, for the chronic session with no electrode insertion there was no overall difference in response between the WT and J20-AD animals. However, for the acute session significant differences between the groups were observed with the J20-AD responses always lower than the corresponding WT response at the same time point (Figure 4).

**Figure 3:**
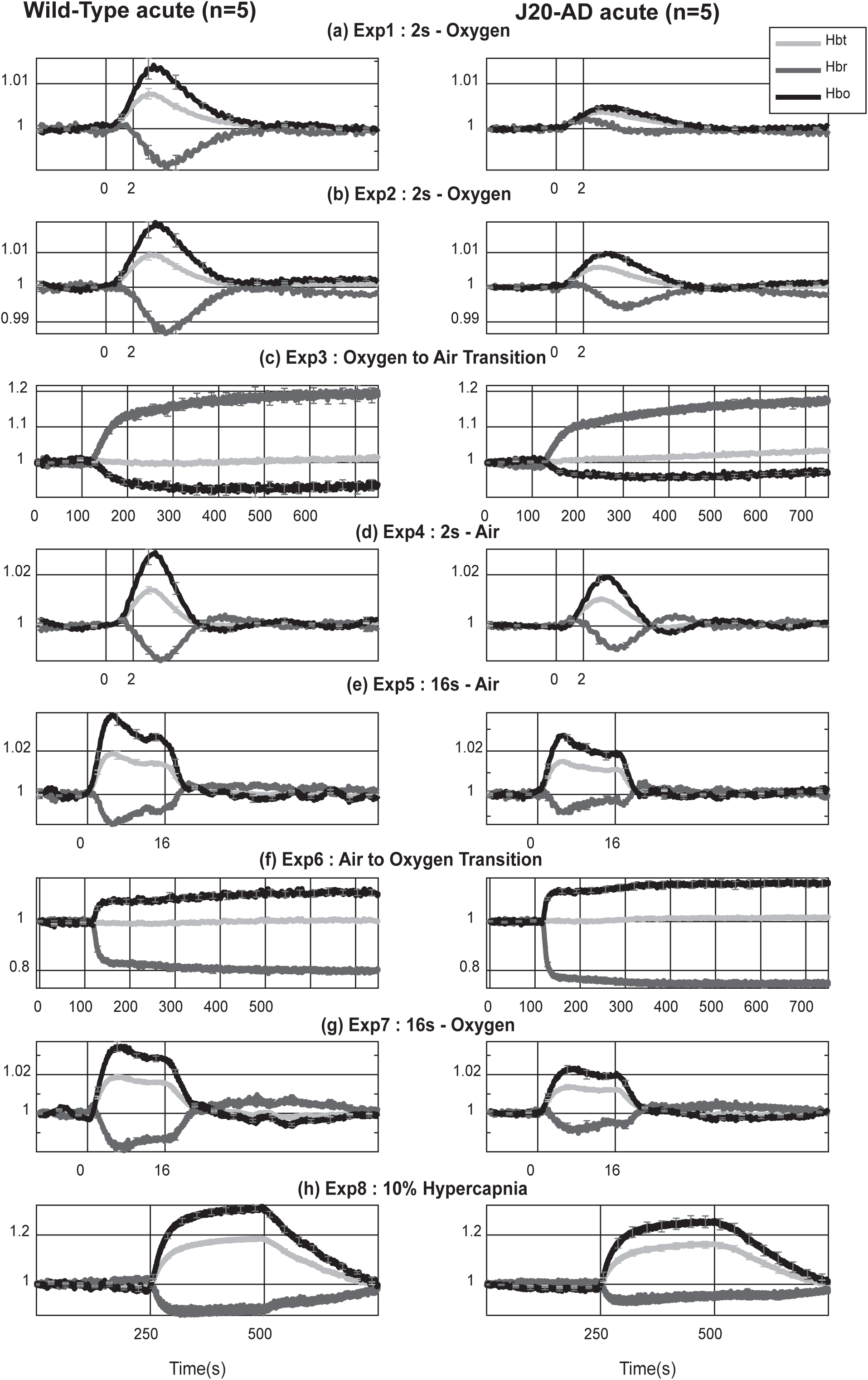
Average hemodynamic time series responses take across all 8 experiments (a-h) in for the acute imaging session with WT in the left column and J20-AD in the right column.

### Concatenated Experimental time course

To better understand the differences between the chronic and acute session results because all our data was collected continuously with small gaps between each of the 8 experiments, we were able to concatenate the data for all experiments to produce a longitudinal record of each session. For example, Figure 5 illustrates the averaged haemodynamic responses to all 8 experiments of the WT group averaged over the first 3 chronic imaging sessions. This figure shows the stability of our chronic anaesthetised protocol insofar as, apart from the expected large shifts caused by the gas challenges (Exp3 and Exp6) and hypercapnia (Exp8), the baseline hemodynamic values were remarkably stable. For instance, individual responses to mechanical whisker stimulation can be seen clearly (Figure 5 insets). Using the same session-long analysis to compare our chronic and acute sessions (Figure 6), it is clear that the baseline values for Hbt, Hbo and Hbr in the chronic conditions for both the WT (black time series) and J20-AD (blue time series) were similarly stable. However, something was clearly different in the acute session in which the recording electrode was introduced prior to collecting the imaging data. In these circumstances the baseline values for both WT and J20-AD seemed to be recovering from a prior event. This was especially evident during the initial phase of the session (exp1 and 2). Also obvious was that the prior event caused the J20-AD perturbation from baseline to be larger than the WT animals. Our supposition is that the perturbation at the beginning of the experimental session was indicative of a cortical spreading depression (CSD) that was caused by the acute electrode insertion. We will discuss this further below, after having presented the electrophysiological and hypercapnia data from our study.

**Figure 5:**
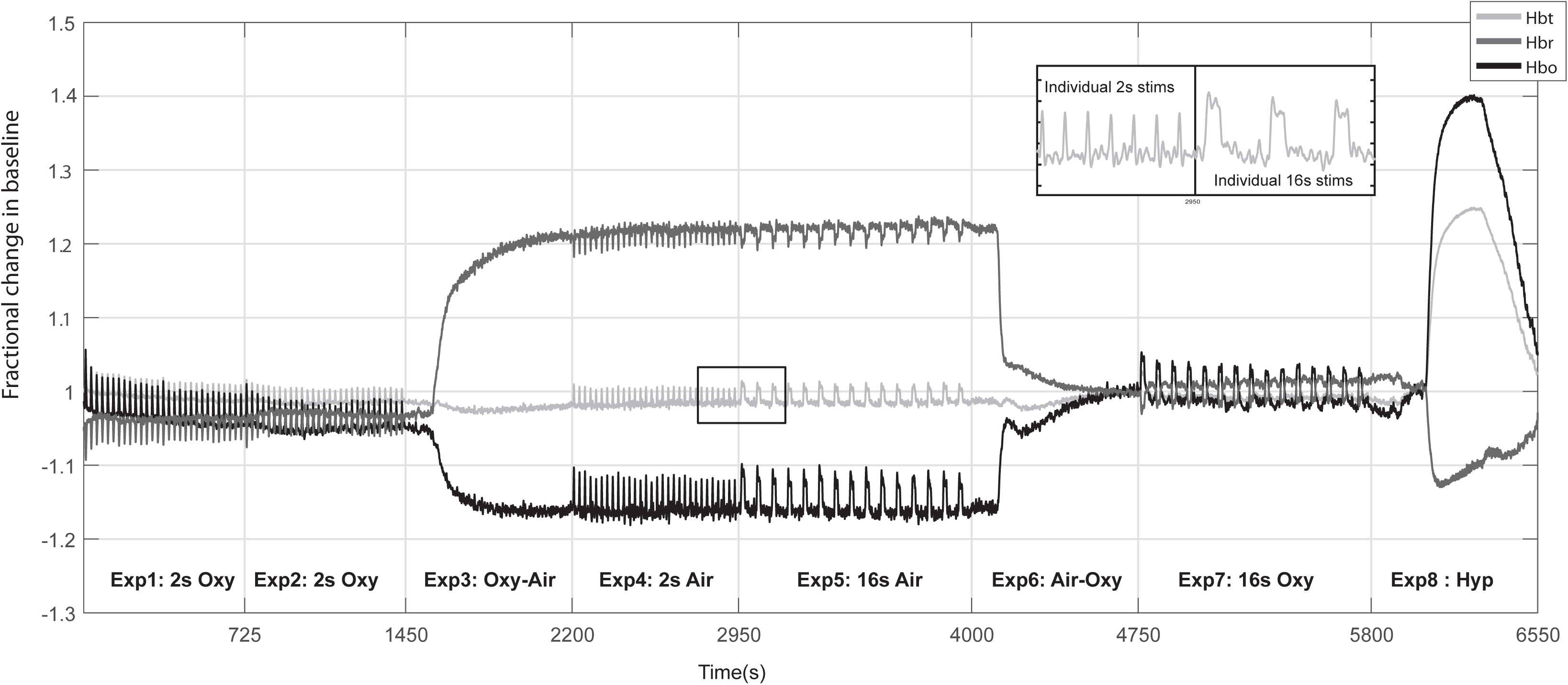
Concatenated hemodynamic times series of responses across experiments for all chronic imaging sessions in the WT imaging group (average of 15 sessions). Hbt= total blood volume, Hbo=Oxyhemoglobin, Hbr=deoxyhemoglobin.

**Figure 6:**
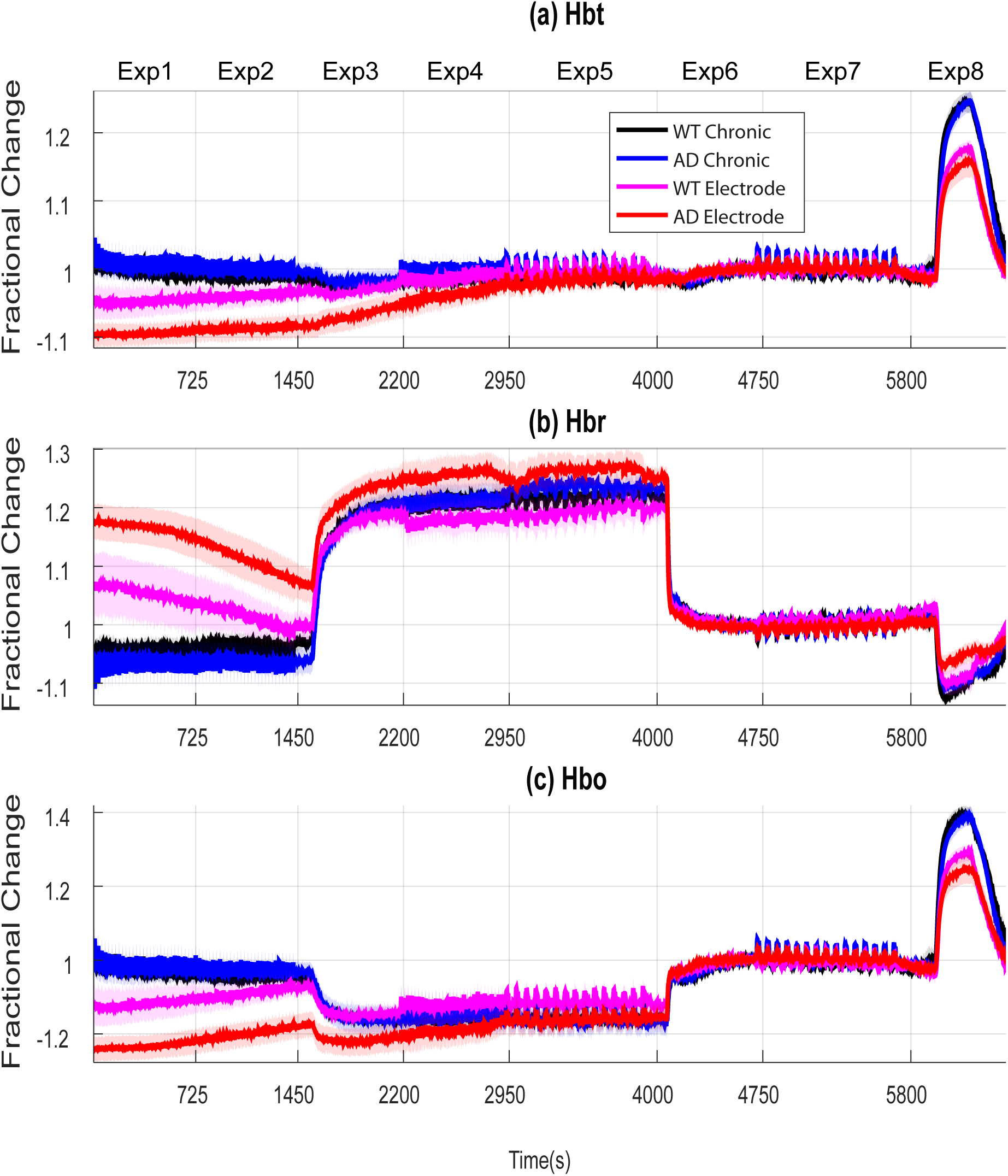
Concatenated hemodynamic time series for chronic and acute WT and J20-AD mice. (a) Hbt (b) Hbr and (c) Hbo.

### Neural Responses to stimulation

The field potential responses from the sensory stimulation experiments (Exps 1,2,4,5, and 7) showed remarkably little difference between the WT and AD groups (Figure 7). A repeated measures ANOVA performed on the averaged evoked field potential response recorded from the middle cortical layers (Figure 7, right hand column), revealed no reliable differences between the animal type (F=2.44, p=0.13), experimental condition (F=1.5, p=0.2) or interaction (F=1.36, p=0.26). The discrepancy between the acute neural and hemodynamic responses is an important issue and will be discussed further below.

**Figure 7:**
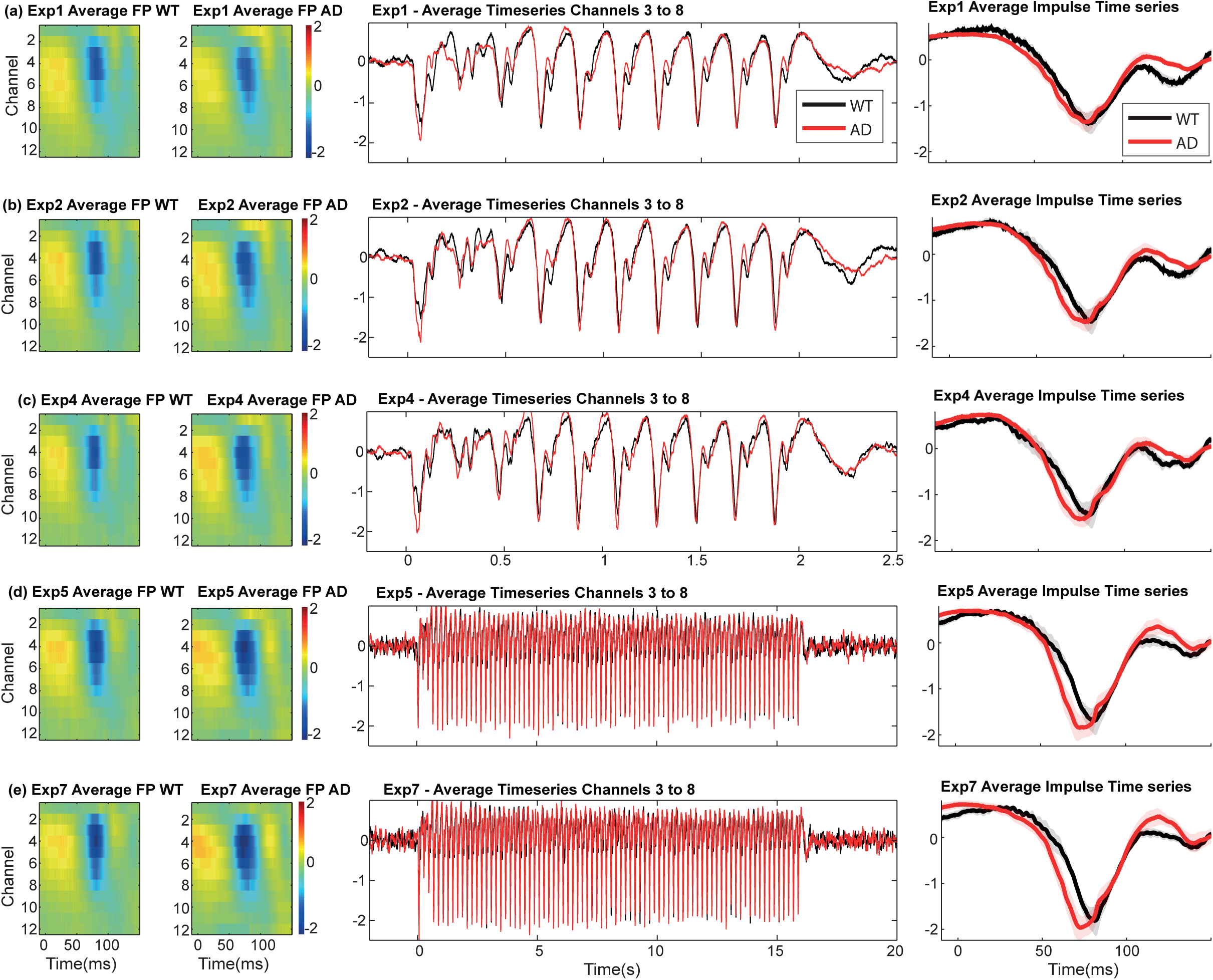
Neural responses from WT and J20-AD mice from stimulation experiments. Left hand images represent the averaged single impulse response in the different stimulation experiments. Middle time series represents the average field potential response from channels 3-8 for WT and J20-AD mice. Right hand column represents the averaged time series impulse response from channels 3 to 8.

### Hypercapnia

To assess the hypercapnia responses recorded during the three chronic imaging sessions were averaged and the mean magnitude of response taken during 250-500s taken. T-tests (two tailed distribution, two-sample equal variance) were conducted to test for significant differences between J20-AD and WT. None of the data for Hbt (p=0.39), Hbo(p=0.28) and Hbr(p=0.21) for the two groups were reliably different. There were also no significant differences between the groups in the final acute imaging session (Hbt, p=0.39, Hbo, p=0.23 and Hbr, p=0.08).

### Compartmental responses and differences between hemodynamic responses evoked by 2s sensory stimulation under air and oxygen

As we have collected 2-dimensional images over time across all our experimental sessions, principal component analysis (PCA) was used to extract responses strongly biased towards the surface vessels as they account for most of the variance in the data (Figure 8). For Hbt the first PCA almost always highlighted major branches of the middle cerebral artery (MCA) responding to the stimulus (Fig 8b). This reflected strong activation of the arterial tree on the surface of the brain. For Hbr the first PCA showed how strongly the surface veins were changing in saturation as Hbr was decreased due to the large increase in fresh blood arriving to the cortex. After selecting arterial and vein regions (Fig 8f-i), clear differences were observed in both groups, not only in the magnitude of the chronic and acute responses (described above) but in the return to baseline depending on whether the animal was breathing 100% oxygen or medical air. Again in both groups under the air condition there was a strong post-stimulation undershoot which was largest in the arterial compartment (Figure 8f). In the pure oxygen condition this overshoot was absent again the importance of this observation will be discussed below.

## Discussion

Our main finding was that differences between the haemodynamic performance of J20-AD mice and WT controls were dependent on experimental procedure. Under stable conditions of chronic imaging, minimal differences between the J20-AD and WT were observed to a wide variety of sensory stimulation and gas challenges. This failed to confirm to previous reports of serious impairments of neurovascular function in the J20-AD model (Lacoste et al., 2013, Ongali et al., 2014, Royea et al., 2017). However, in our final experiment, which involved acute experimental procedures a clear impairment of haemodynamic responses in the J20-AD animals was observed, similar to previous reports (Lacoste et al., 2013, Ongali et al., 2014, Royea et al., 2017). However, compared to the chronic imaging sessions, we found that the magnitude of evoked-responses of both WT and J20-AD mice were reduced, with the J20-AD mice affected to a significantly greater extent. We draw a general conclusion that well-controlled procedures are necessary to separate the effects of methodological confounds from genuine effects of pathophysiological changes induced by the disease state being tested.

### Preservation of neurovascular coupling in the J20 mouse model of Alzheimer’s disease

Why in the stable chronic recording phase of our study did we fail to replicate the neurovascular deficits reported previously? First, we used similar whisker stimulation protocols to the studies that reported 35-50% reductions in the cerebral blood flow responses of J20-AD mice from 7 months of age. Our subjects were 9-12 months old at the start of the study. Both our and previous experiments were all performed with anaesthetised mice. However, previous studies (Lacoste et al., 2013, Ongali et al., 2014, Royea et al., 2017) conducted few stimulation presentations (4-6 stimulation trials per animal), in the anaesthetised mouse immediately following acute procedures to thin the skull. At the outset, we sought to extend these findings by using 2D-OIS to resolve changes in different vascular compartments with repeated testing in a chronically prepared preparation. Informed by our previous work with an awake head restrained preparation (Sharp et al., 2015), we adopted a refined anaesthetised protocol that permitted extended (3hr) imaging sessions repeatedly over weeks. Under anaesthesia, the haemodynamic responses evoked by sensory stimulation in this chronic preparation were comparable to those seen previously with awake animals (Sharp et al., 2015). Another difference was the present protocol involved a more detailed set of experimental variables, i.e. sequences of multiple trials (120) with different duration whisker stimulations (2s and 16s), with and without vascular reactivity gas challenges. Also, our protocol enabled us to generate an average experimental session by averaging the sequence of hemodynamic time series from all chronically prepared subjects (Figure 5 and 6). These data show, not only that responses to individual sensory stimulations could be observed, but that the haemodynamic baseline values were stable throughout the session. Under our more stringent chronic conditions and more detailed testing we failed to replicate the previously reported haemodynamic deficits in J20-AD mice. Therefore, our conclusion must be that neurovascular function is still largely preserved in the J20-AD mouse in this age range.

### Acute experimental conditions

Given these essentially negative results from the first three chronic experimental sessions, we looked for correspondences between haemodynamic and neural responses in a final acute experimental session. Significantly, this phase of the study involved acute experimental conditions akin to those used in the previous investigations (Lacoste et al., 2013, Ongali et al., 2014, Royea et al., 2017). After the electrode insertion we noted that that baseline hemodynamics were altered. Specifically, there was a significant reduction in blood volume and saturation that gradually recovered to normal stable levels towards the end of the session. Significantly, the baseline changes were larger for the AD mice compared with the WT controls. We therefore suggest that acute procedures, in our case the insertion of a multi-channel electrode, likely causes a cortical spreading depression (CSD). This is characterised by an alternating vasoconstriction and dilation that can last over 90 minutes, and is known to affect neurovascular coupling profoundly (Piilgaard and Lauritzen, 2009, Chang et al., 2010). Therefore, differential rates of recovery from CSD between the J20-AD and WT mice is the most likely explanation of the differences in haemodynamic performance between the groups. Also important, Chang et al (2010) reported that hemodynamic recovery from CSD takes longer than the recovery of neural activity. This could explain the lack of correspondence between the neural (no differences) and haemodynamic measures (significant differences) between the WT and J20-AD groups in the final acute phase of our study. Supporting this interpretation, electrode insertion in mice induced a robust CSD response across the whole cortex (Eles et al., 2018). If CSD is the culprit, why should J20 mice be more susceptible to its deleterious effects? It is known that J20 mice exhibit hypertrophic astrocytes and activated microglia in the cortex near beta amyloid plaques (Ameen-Ali et al., 2017). If these cells were sensitized as part of the disease process, the CSD event could be more profound in J20-AD animals. The larger baseline disturbance in the J20-AD animals at the start of recording in the final acute experimental session would be consistent with this suggestion (Figure 6).

### Oxygen Vs Air response shape

A further unexpected result was that, regardless of mouse model, the shape of hemodynamic response changed according to whether the animal was breathing oxygen or air. The overall response magnitude was reduced and there was an undershoot in the return to baseline when the animals were breathing air, compared with when they were breathing oxygen. This difference was driven by the arterial response (see Figure 8). In contemporary neurovascular research the prevailing view is that coupling is more likely to be neurogenic in origin, compared to metabolically driven changes in blood flow (Kennerley et al., 2012, Urban et al., 2012). However, as the neural responses evoked by 2s sensory stimulation were remarkably similar between the air and oxygen conditions (Figure 7), the observed differences appear to be a direct effect of inspired oxygen controlling the arterial response, particularly in the return to baseline. Previous *in vitro* experiments (Gordon et al., 2008), reported that levels of oxygen can dictate whether an artery dilates or constricts in response to a stimulus, and that this is mediated by different responses of astrocytes. Interestingly, a recent 2-photon microscopy study (Uhlirova et al., 2016) reported similar arterial responses to mouse sensory stimulation. In some cases the responses had a bi-phasic shape (similar to our air condition) while in others, there was have a gradual return to baseline without overshoot (similar to our oxygen condition). No reasons were given to why the same sensory stimulus produced a varying arteriolar responses. However, an important conclusion from this aspect of our study is that tissue oxygen levels may be an important determinant of neurovascular control and warrants further investigation.

### Limitations of the study

Our current spectroscopy algorithm requires an estimation of both a level of blood volume and saturation. Although, previous published research used LDF, which has a similar weakness, arterial spin labelling studies suggest that baseline CBF is reduced in J20-AD mice (Hebert et al., 2013). We mitigated these effects by measuring changes in baseline volume and saturation in our animals when switching from breathing oxygen to air. If there were differences in saturation between the two groups, this would be evidence that our estimated baselines values were inaccurate. However, the consistency of the results between the WT and AD mice suggests that under the oxygen condition, our estimates were appropriate. Moreover, even if estimates of volume and saturation were wrong, the fractional change in Hbt response would be unaffected. We have run the data across a range of assumed Hbt baselines and this produces no difference in the fractional change in Hbt response (Figure 9).

**Figure 9:**
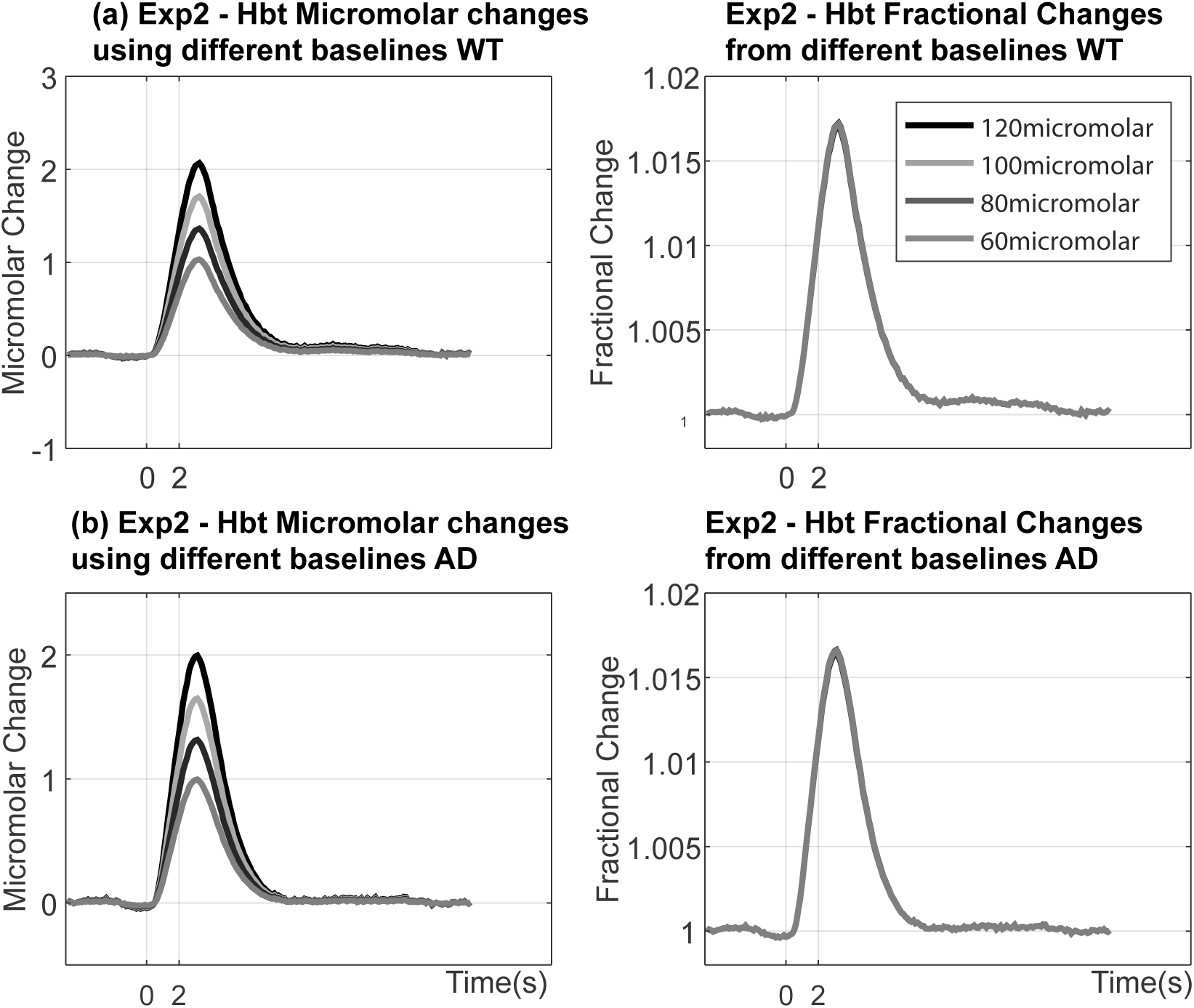
Effect on varying baseline blood volume concentration on fractional hemodynamic responses for WT and J20-AD mice. (a) Left hand side micromolar responses using varying baseline WT mice. Right hand side corresponding fractional changes from the varying Hbt baselines. (b) Left hand side micromolar responses using varying baseline J20-AD mice. Right hand side corresponding fractional changes from the varying Hbt baselines.

Finally, it must be noted that all our experiments were performed under anaesthesia, which of course is a non-physiological state. To some extent this is mitigated by our previous investigation (Sharp et al., 2015) that showed the responses in the present study were comparable to those seen previously with awake animals. On the positive side, the current anaesthetised preparation provides a more stable imaging platform where the effects of controlled small mechanical stimulations can be observed. Importantly, baseline perturbations, such as caused by the insertion of an electrode, are easier to detect in the absence of uncontrolled motivational and movement variables that are often inherent in awake experimental protocols.

## Conclusions

Contrary to previous reports of neurovascular deficits J20-AD mice, under the stable chronic imaging conditions we found that the neurovascular performance of J20-AD did not differ reliably from that of WT controls. However, the acute phase of the current study showed the J20-AD mice to be more susceptible to disruptive interventions, such as the acute experimental procedures used in the current and previous investigations. The J20-AD mouse model may, therefore, represent an ideal model to explore the effects of mixed pathologies with conditions such as atherosclerosis that would further compromise neurovascular function.

## Acknowledgements

This work was funded by Alzheimer’s Research UK (ARUK-IRG2014) and the Medical Research Council UK (MR/M013553/1). The authors would like to thank Michael Port for his technical assistance.

